# Fused in sarcoma undergoes cold denaturation: Implications on phase separation

**DOI:** 10.1101/2022.02.08.479505

**Authors:** Sara S. Félix, Douglas V. Laurents, Javier Oroz, Eurico J. Cabrita

**Affiliations:** Associate Laboratory i4HB - Institute for Health and Bioeconomy, NOVA School of Science and Technology, Universidade NOVA de Lisboa, 2819-516 Caparica, Portugal; UCIBIO, Department of Chemistry, NOVA School of Science and Technology, Universidade NOVA de Lisboa, 2819-516 Caparica, Portugal; Instituto de Química Física “Rocasolano”, Consejo Superior de Investigaciones Científicas, c/ Serrano 119, Madrid 28006, Spain

**Keywords:** Cold denaturation, Fused in Sarcoma, Liquid-liquid phase separation, NMR

## Abstract

The mediation of fused in sarcoma (FUS) protein liquid-liquid phase separation (LLPS) is generally attributed to the low-complexity and disordered domains, while the role of its folded domains remains unknown. In this work we questioned the role of the folded domains on the full-length (FL) FUS LLPS and studied the influence of several metabolites, ions and overall conditions on the LLPS process using turbidity assays, differential interference contrast microscopy and nuclear magnetic resonance spectroscopy. We demonstrate that FL FUS LLPS is highly responsive to the surrounding conditions, and that overall intrinsic disorder is crucial for LLPS. To promote such disorder, we reveal that the FUS RNA-recognition domain (RRM) and the zinc-finger motif (ZnF) undergo cold denaturation above 0ºC, at a temperature that is determined by the conformational stability of the ZnF domain. We hypothesize that, in cold shock conditions, cold denaturation might provide a pathway that exposes additional residues to promote FUS self-assembly. Such findings mark the first evidence that FUS globular domains may have an active role in stress granule formation in cold stress.

## Introduction

Liquid-liquid phase separation (LLPS) of proteins is a recently recognized biological mechanism that describes the formation of membraneless compartmentalization in the cellular context (Brangwynne *et al*, 2009; Elbaum-Garfinkle *et al*, 2015; Brangwynne, 2013; Banani *et al*, 2017). The biomolecular condensates assembled by LLPS are typically comprised of RNA and complex proteins, and have been shown to behave as fluid droplets with liquid-like properties (Buchan & Parker, 2009; Protter & Parker, 2016).

In the past years, mounting evidence has been collected showing that biomolecular condensates rule several biochemical processes, ranging from nucleic acid processing to age-related neurodegeneration (Li *et al*, 2013; Kwiatkowski *et al*, 2009; Sun & Chakrabartty, 2017). Among the important types of biomolecular condensates are the stress granules that assemble from pools of untranslated mRNA, RNA-binding proteins (RBPs) and factors involved in translational repression and mRNA decay, during stress conditions. In the cytoplasm, stress granules regulate RNA metabolism, stabilize translation preinitiation complexes, and modulate signaling pathways in response to stress stimuli (Courchaine *et al*, 2016; Jain *et al*, 2016; Buchan, 2014).

Fused in sarcoma (FUS) is a complex prion-like RBP that is found in stress granules (Bentmann *et al*, 2012; Bosco *et al*, 2010; Gal *et al*, 2011). This protein is predominantly localized in the nucleus and is involved in the transcriptional regulation function of cells (Chen *et al*, 2019). However, FUS can shuttle between the nucleus and cytoplasm, where it can be recruited into stress granules (Gal *et al*, 2011; Sama *et al*, 2013). The array of broad functions of FUS lies in the multidomain character of this protein. FUS contains a N-terminal low complexity (LC) domain that is enriched with glutamine, glycine, serine and tyrosine (QGSY) residues, known to have an important role on protein self-association through the mediation of LLPS (Fig. 1a) (Burke *et al*, 2015; Murakami *et al*, 2015). Also, the protein contains a nuclear export signal (NES) and a proline-tyrosine nuclear localization signal (PY-NLS), involved in the nuclear-cytoplasmatic shuttling. Mediation of protein-nucleic acid interactions is governed by the three arginine-glycine-glycine (RGG) boxes and, by the structured RNA recognition motif (RRM) and the cysteine_2_-cysteine_2_ zinc finger (ZnF) domain (Deng *et al*, 2014; Yang *et al*, 2010; Loughlin *et al*, 2019). Apart from the RRM and ZnF motifs, the remaining domains are predicted to be highly disordered (Rogelj *et al*, 2012).

**Figure 1.**
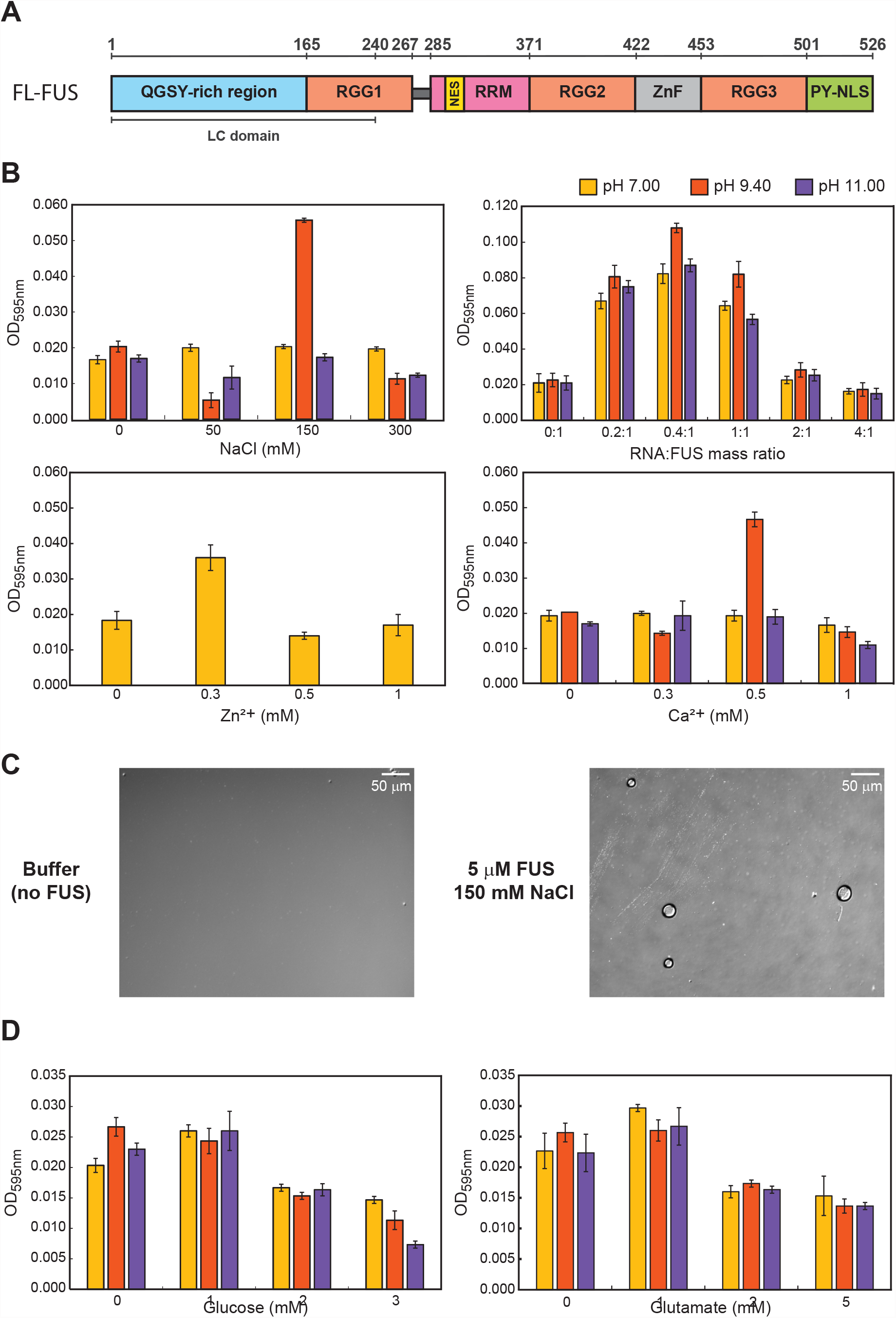
FL FUS phase separation is highly sensitive to the sample conditions. (A) FL FUS domain architecture.; (B) Liquid-liquid phase separation of 5 μM FL FUS increases in the presence of specific concentration of NaCl (150 mM), RNA:FUS mass ratio (0.4:1), Zn^2+^ (0.3 mM) and Ca^2+^ (0.5 mM). This promotion of LLPS is not proportional to metabolite concentration, indicating the tight regulation of the phase separation of FUS. Overall, the phase separation is maximized when the pH=pI of FUS. Data are represented as triplicate mean ± SD.; (C) DIC microscopy images of 5 μM FUS in 150 mM NaCl shows liquid-liquid phase separation droplets.; (D) Liquid-liquid phase separation of 5 μM FL FUS decreases in the presence of stabilizing metabolites (glucose and glutamate). Color code in (B) is maintained in (D).

FUS is able to undergo LLPS both *in vivo* and *in vitro*, self-assembling into liquid-like stress structures at concentrations in the low micromolar range (Burke *et al*, 2015; Patel *et al*, 2015; Wang *et al*, 2018). FUS stress granules are highly dynamic structures, displaying constant exchange of components with the cytoplasm and rapid internal rearrangement (Patel *et al*, 2015). Since the cell uses dynamic liquid compartments to perform physiological relevant functions, there are mechanisms that constantly control the thermodynamically driven aggregation processes. Hence, FUS-related diseases are often associated with late onset pathology, when the machineries responsible for the maintenance of quality control in the cells start to degenerate (Shin & Brangwynne, 2017; Patel *et al*, 2015). Stress granules often accumulate misfolded proteins that can evolve into aberrant aggregated states (Boeynaems *et al*, 2017; Mateju *et al*, 2017). This stimulates the protein quality control (PQC) machinery to produce factors that promote condensate dissolution or degradation (Ganassi *et al*, 2016). As ageing occurs, there is a progressive loss of proteostasis mainly due to loss-of-function mutations in PQC factors. This leads to the increase of misfolded proteins and hampered clearance of aberrant condensates with age (Alberti & Hyman, 2021). Aberrant aggregated forms of FUS are a major component of pathological inclusions found in 5% of all forms of amyotrophic lateral sclerosis (ALS) and in 8% of all cases of frontotemporal lobar degeneration (FTLD) (Buchan & Parker, 2009; Yang *et al*, 2010).

FUS plays an imperative role in cellular stress response. Cells exposed to environmental stress, such as oxidative stress, hypoxia, viral infections or osmotic stress, actively promote cytoplasmatic FUS stress granule formation (Bentmann *et al*, 2012; Dormann *et al*, 2010; Bosco *et al*, 2010). Moreover, interestingly, FUS phase separation has been shown to be highly enhanced at low temperature, suggesting that cold stress can also induce the stress granule assembly (Burke *et al*, 2015). In fact, cold stress has been identified as a trigger for stress granule assembly in yeast and mammals, acting as a protective mechanism against cell death during hypothermia, which suggest that FUS might have an active role in cold shock response (Al-Fageeh & Smales, 2006; Hofmann *et al*, 2012).

In recent years there have been vast developments in the LLPS research area; from the sequence determinants of protein LLPS to the favorable molecular interactions (Dignon *et al*, 2018; Wang *et al*, 2018; Mitrea & Kriwacki, 2016). However, there is still a need to investigate the specific triggers and determinants that lead to LLPS and eventually to the transition to proteinaceous deposits (Sama *et al*, 2014).

Since the LLPS process relies on the establishment of weak interaction networks, the conviction is that different environmental conditions and stresses, and consequently different interactions, should uniquely influence FUS and the LLPS mechanism (Uversky *et al*, 2015; Darling *et al*, 2018; Protter & Parker, 2016). Remarkably, current studies are mainly based on fragmented FUS domains. Low-complexity domains of RNA-binding proteins, such as FUS, are often sufficient for phase separation (Burke *et al*, 2015; Molliex *et al*, 2015; Xiang *et al*, 2015). However, studies of full-length (FL) FUS and the interplay between disordered and folded domains are still scarce, which is possibly due to the elevated tendency for the FL FUS to stick to surfaces and to readily aggregate (Emmanouilidis *et al*, 2021; Wang *et al*, 2018).

In this work, to address the lack of insight on the phase separation of FL FUS and how the synergism among the different domains impact the mechanism, the research presented is solely based on FL FUS. We use a novel sample preparation and handling protocol to avoid surface binding and aggregation. We seek to unravel the connection between cold stress and FUS phase separation and to assess the influence of diverse environmental conditions in the formation of FL FUS stress granules, using a combination of nuclear magnetic resonance (NMR) spectroscopy, differential interference contrast (DIC) microscopy and turbidity assays.

Insights on the role of the environment on the LLPS mechanism can prove valuable in understanding the physiological and pathological conditions that trigger FL FUS stress granule assembly. Our results provide the first structural insight into the influence of cold stress on the FUS protein and how phase separation can be regulated in cold stress conditions.

## Results

### Charged metabolites and preferred global neutral charge enhance FUS phase separation

Charged metabolites are ubiquitous in cells, being crucial for almost all cellular processes. For this reason, the influence of inorganic salt ions (NaCl), metal ions (Ca^2+^ and Zn^2+^) and RNA in the FUS LLPS mechanism was tested by turbidity assays, following the OD at 595 nm. It is important to mention that despite turbidity assays of FL FUS have been previously reported, in most cases the assays are done with MBP-FUS in the presence of TEV protease (Burke *et al*, 2015; Ahlers *et al*, 2021; Kaur *et al*, 2019). Since it is known that FUS phase separation is highly sensitive to crowding and prone to unspecific interactions with proteins, it is possible that the presence of cleaved MBP and TEV protease in the solution could affect the turbidity results and analysis (Kaur *et al*, 2019; Kang *et al*, 2019; Lin *et al*, 2015). For this reason, we performed the LLPS turbidity assays with pure non-tagged, FL FUS (Fig. 1a).

FL FUS phase separation displayed a high sensitivity to the nature and concentration of each metabolite in a pH-dependent manner (Fig. 1b). LLPS was observed to be maximal when the protein’s overall net charge is zero, at pH 9.40. Among all tested metabolites, FUS LLPS was notably promoted in the presence of 150 mM NaCl. Similar results have been previously observed with a highly basic protein, CAPRIN1, that undergoes phase separation in the presence of negatively charged phosphate buffer, suggesting that charge screening might be crucial for LLPS promotion (Wong *et al*, 2020). FUS phase separation was further confirmed by DIC imaging choosing the condition of 150 mM NaCl (Fig. 1c).

The modulation of FUS phase separation by the presence of metal ions was however different. The behavior of divalent transition metals and divalent alkali earth cations such as Zn^2+^ and Ca^2+^ is distinct from that of monovalent ions such as Na^+^. Such metal ions can form coordination complexes leading to stronger interactions. FUS contains a zinc-finger motif in which a Zn^2+^ cation is coordinated. In both tested cases, the metal ions induced FUS phase separation at concentration about 300-fold lower compared to the tested monovalent ions in order to reach a similar degree of turbidity (Fig. 1b).

### Stabilizing metabolites hinder FUS liquid-liquid phase separation

Given that both glucose and glutamate are two of the most abundant metabolites found in cells, and that both are known to act as stabilizing metabolites by inducing preferential hydration of proteins, we investigated how these of metabolites would affect FUS LLPS (Psychogios *et al*, 2011; Scimemi & Beato, 2009).

Our results showed that in the presence of increasing glucose or glutamate concentrations, FUS phase separation was inhibited at all pH values studied (Fig. 1d). By inducing preferential hydration, glucose and glutamate are capable of displacing water molecules from the protein surface leading to structural compaction by reducing the protein total exposed surface area (Arakawa & Timasheff, 1982, 1984). The observed decrease of FUS phase separation by increasing concentrations of glucose and glutamate suggests that the intrinsic disorder of FUS might be important for the phase separation process, as the reduced surface area might lead to inaccessibility of FUS phase-promoting interacting sites.

### FUS undergoes cold denaturation which depends on the ZnF stability

To understand how cold shock could affect FUS phase separation, our first approach was to investigate the impact of cold stress on FUS structure by NMR spectroscopy. By the acquisition of ^1^H-^15^N TROSY-HSQC and ^1^H-^15^N HSQC spectra, we followed FUS structural changes induced by temperature, from physiological temperature (37ºC) to mild hypothermia (35ºC-25ºC) and to severe hypothermia (15ºC-5ºC) in the absence of ZnCl_2_.

As the temperature decreases, the signals from the folded regions (corresponding to the RRM and ZnF domain) lose intensity and re-emerge in the disordered region of the spectrum, between 8-9 ppm (Fig. 2a). This is especially clear in the 1D ^1^H projection of the ^1^H-^15^N TROSY-HSQC spectra (Fig. 2b) where, as the temperature lowers, the intensity of the proton signals in the unfolded region increases. Using available assignments in the BMRB database (BMRB LC #26672 and RRM #34259), we could identify several isolated cross-peaks belonging to the LC and RRM domains from FUS. We analyzed the intensity of the isolated LC domain and RRM cross-peaks as the temperature decreased (Fig. 3a). While the LC domain cross-peaks disappeared at higher temperature (>30ºC) due to chemical exchange, those belonging to the folded RRM domain disappeared at lower temperature (<15ºC), indicating domain unfolding. When the temperature was increased back to 37ºC, the cross-peaks reverted to the initial position, showing that the process is reversible.

**Figure 2.**
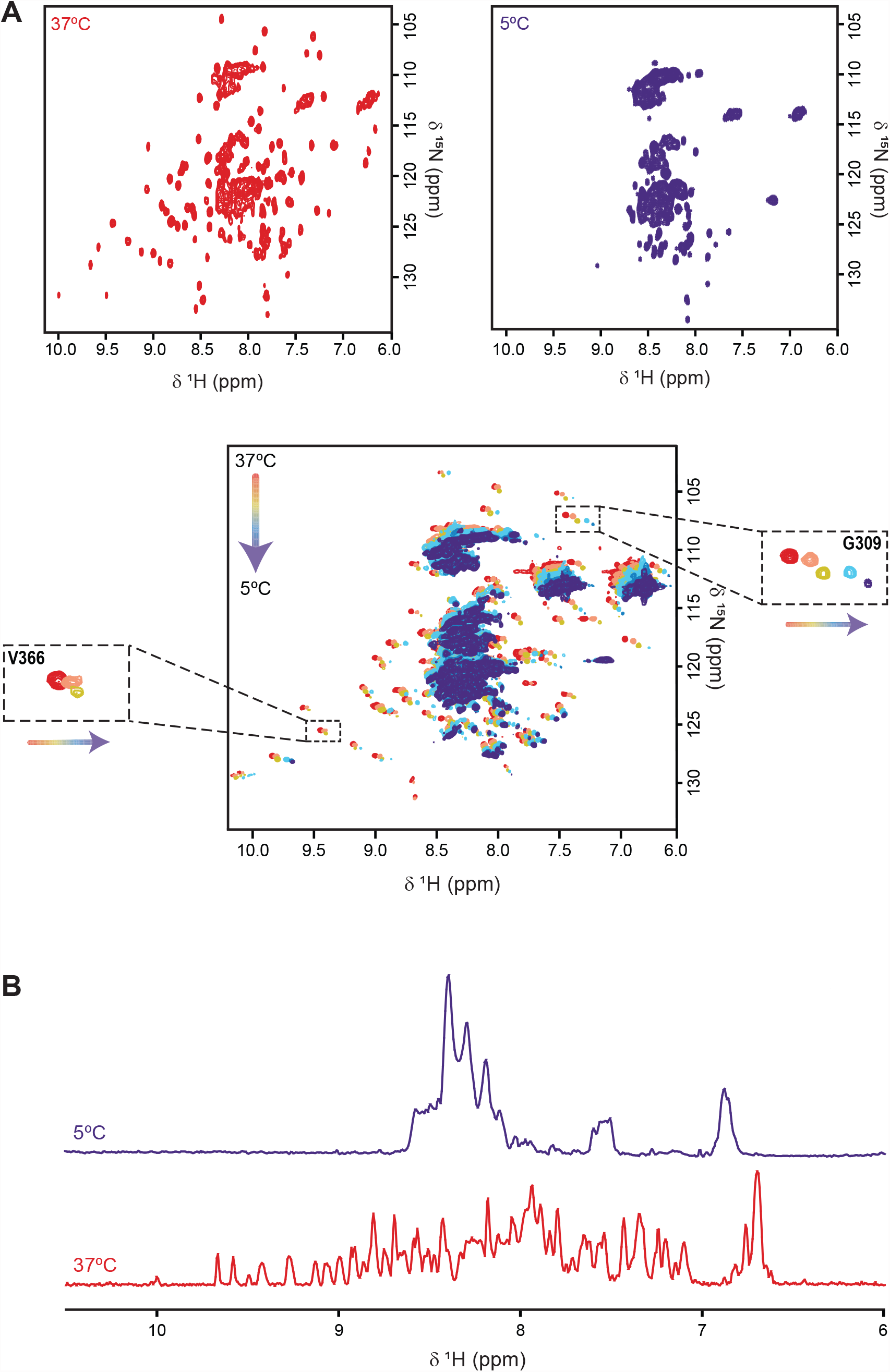
FUS globular domains undergo cold denaturation above 0ºC in the absence of Zn^2+^. (A) ^1^H-^15^N TROSY-HSQC spectra of 30 μM FL FUS from 37ºC to 5ºC revealing cold denaturation. Example of valine 366 from the RRM domain which disappears at low temperature opposed to glycine 309 from disordered RGG2.; (B) ^1^H projections of FUS ^1^H-^15^N TROSY-HSQC spectra at 37ºC and 5ºC showing the collapse of peak chemical shifts in the disordered region 8-9 ppm.

**Figure 3.**
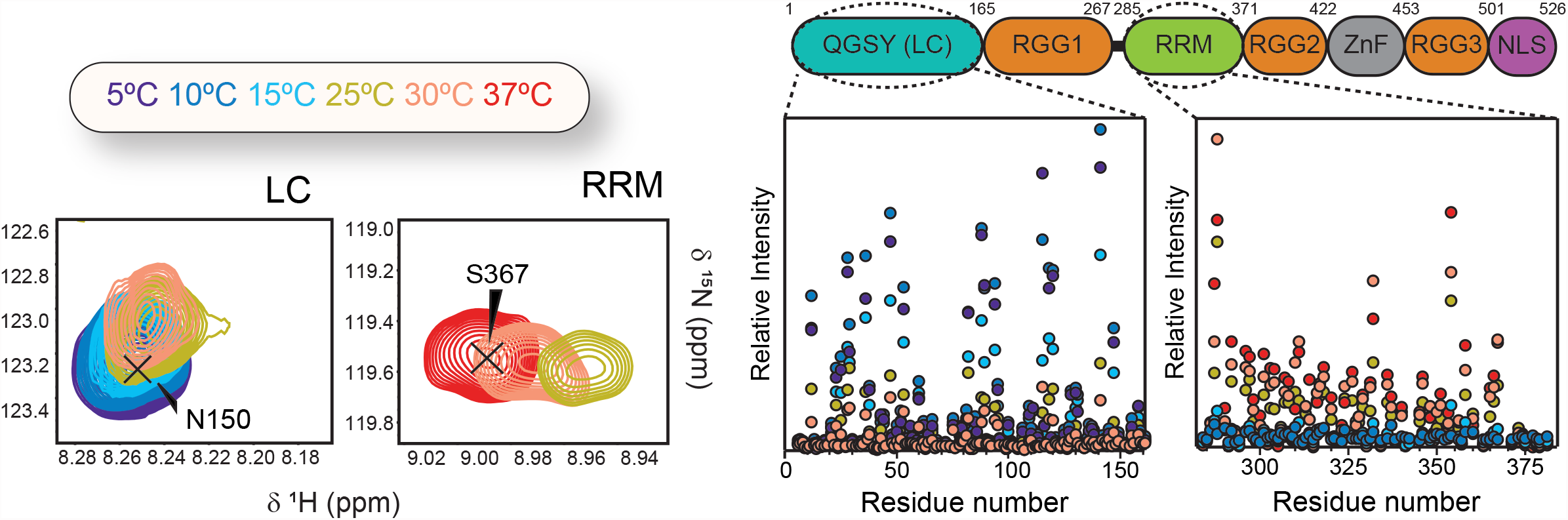
Analysis of the NMR peaks from order vs disordered regions of FUS. Representative regions of the ^1^H-^15^N HSQC spectra at different temperatures (as indicated in the color code) showing isolated cross-peaks corresponding to the LC and RRM domains of FUS (left panel). LC moiety peaks disappears at higher temperature due to chemical exchange, while RRM peaks fade out at low temperature, indicating domain unfolding. Intensity plots (right panel) show the reverse temperature dependance of the NMR signals from the LC and RRM domains.

These results showed that, in the absence of ZnCl_2_, FUS folded domains undergo reversible cold denaturation, which renders an unstable ZnF motif.

### Cold denaturation of FUS is independent of phase separation

To assure that the changes in NMR signal intensity were not due to FUS phase separation, turbidity was checked at each temperature (Table S1) and peak analysis showed that ^1^H linewidths remain almost unchanged at each temperature (Fig. 3b). NMR signal linewidth would be expected to increase proportionally as phase separation occurred due to the higher viscosity of the sample (Burke *et al*, 2015). Hence, during the temperature variation experiment, the sample remained dispersed and the drop in intensity in the folded regions can be attributed to local unfolding as the temperature decreases. Nevertheless, to corroborate that the spectral changes were a result of cold denaturation and not phase separation or a combination of both, the NMR experiments were also performed with FUS in the phase-separated state, and in the presence of 4% 1,6-hexanediol. This aliphatic alcohol has been showed to completely disrupt phase separation of FUS at 4% concentration (w/v) (Kroschwald *et al*, 2017). This experiment was used as a negative control, where FUS is completely dispersed in solution (Table S1). The same behavior was observed in the NMR spectra both in presence and in the absence of 4% 1,6-hexanediol, indicating that the spectral changes do not come from phase separation and can be attributed to cold denaturation (Fig. 4). This is corroborated by NMR experiments of FUS in the phase-separated state that served as a positive control which showed the complete loss of signal upon LLPS (Fig. S1).

**Figure 4.**
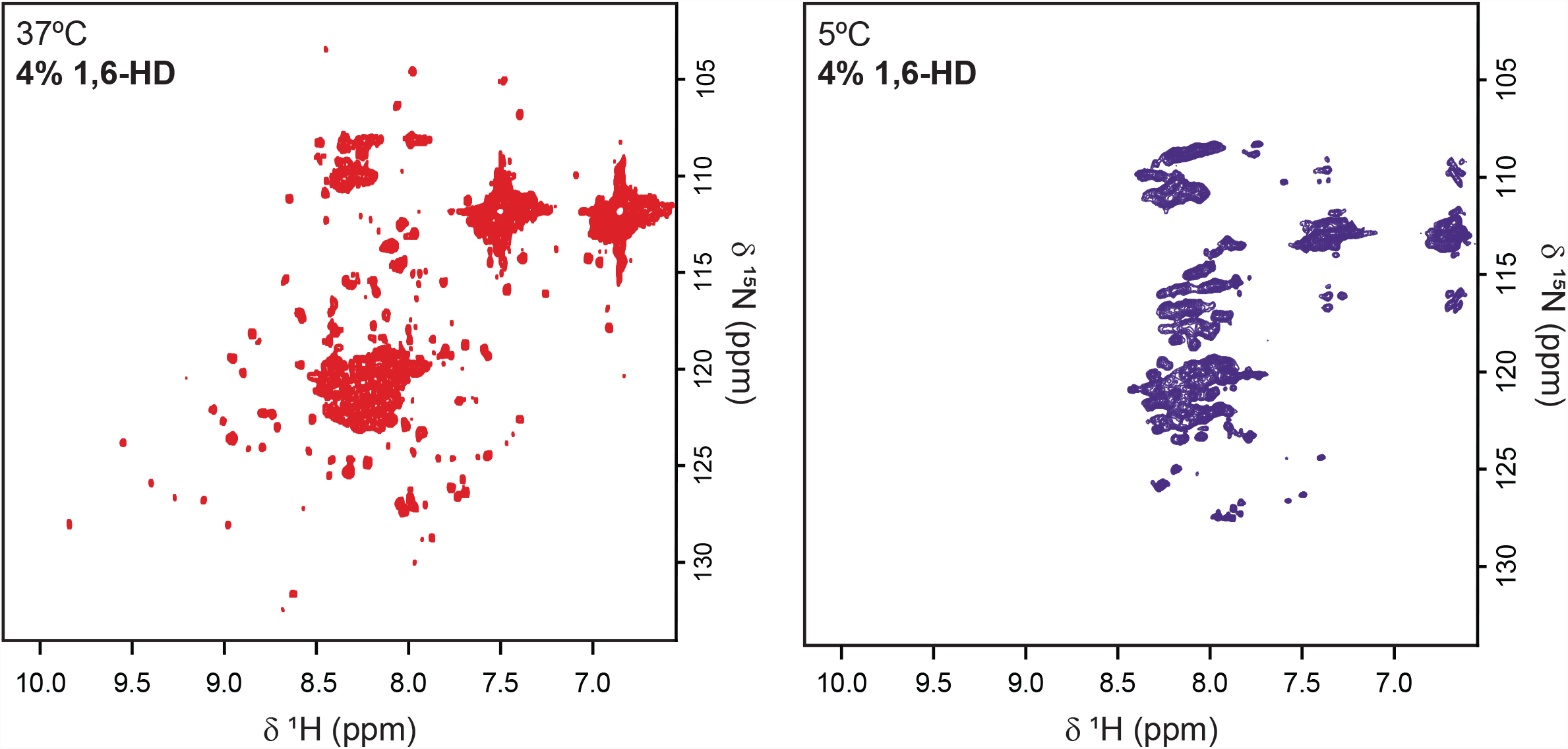
Cold denaturation of FUS is still detected in the presence of liquid-phase inhibitor 1,6-hexanediol. Negative control ^1^H-^15^N HSQC spectrum of 30 μM FL FUS in the presence of 4% 1,6-hexanediol (HD), indicating that the spectral changes are not due to phase separation.

Inspired by the standard procedure to induce cold denaturation of proteins above the water freezing temperature using denaturing agents, urea was added to FUS samples and spectral changes over a temperature range were followed (Privalov, 1990; Wong *et al*, 1996; Agashe & Udgaonkar, 1995). Chaotropic agents such as urea are able to shift cold denaturation equilibrium towards higher temperatures in order to detect unfolded populations. In the presence of 750 mM urea, cold denaturation of FUS is indeed detected at higher temperatures compared to in the absence of urea (Fig. 5). This experiment thus provides additional evidence that folded domains of FUS are undergoing cold denaturation.

**Figure 5.**
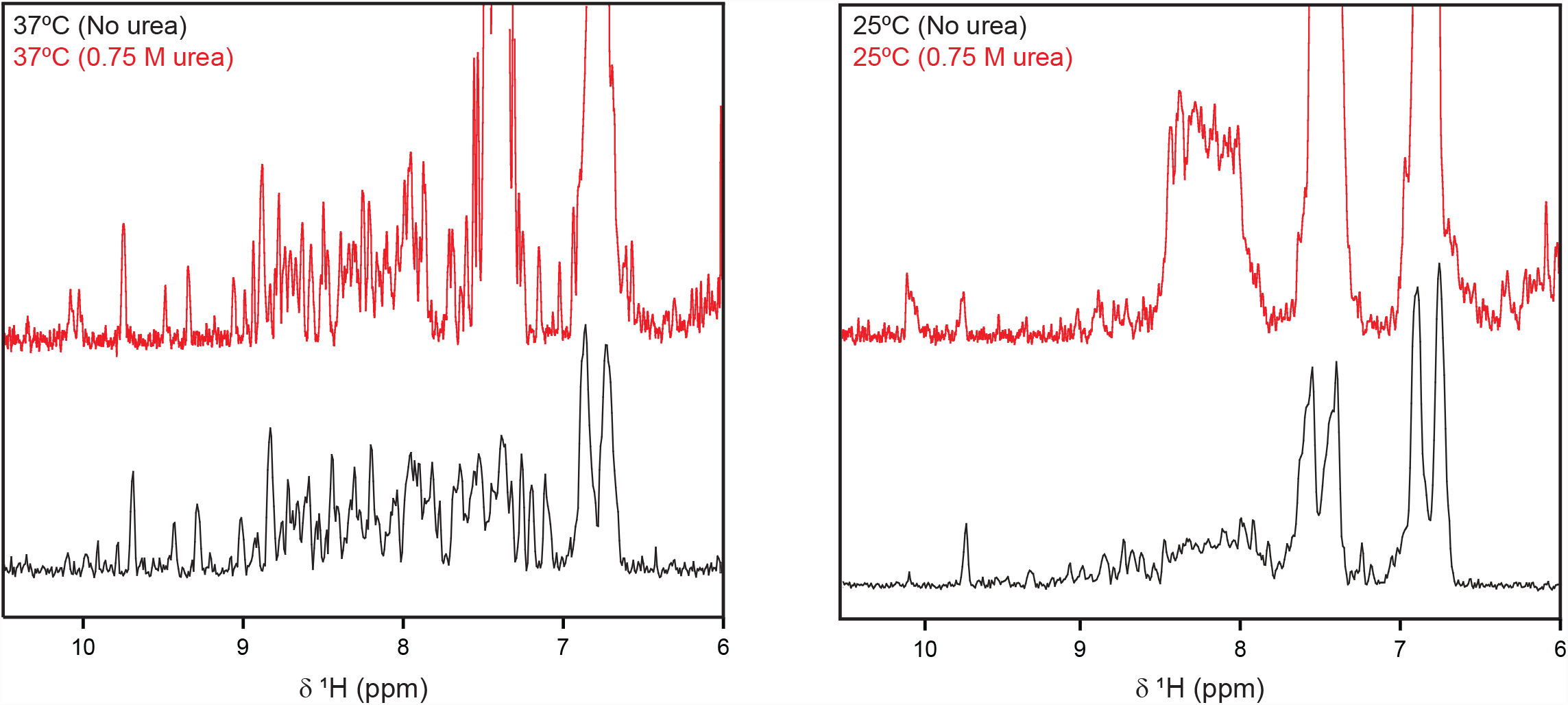
Urea shifts cold denaturation temperature equilibrium towards higher temperatures. ^1^H NMR spectrum reveals that 30 μMFL FUS in the presence of 0.75 mM urea undergoes cold denaturation at higher temperature compared to without urea. In the presence of urea, there is a large increase of intensity in the region 8-9 ppm at 25ºC (right panel) which indicates substantial protein unfolding, whereas no significant changes are seen at 37ºC (left panel). Without urea the unfolding only starts at ≤15ºC.

### The presence of Zn^2+^ protects FUS from cold denaturation

Remarkably, when 100 μM of ZnCl_2_ was added to FUS, cold denaturation was not observed at the same degree when compared to samples without ZnCl_2_ (Fig. 6). In the presence of Zn^2+^, FUS samples are overall more stable, which could be attributed to the native coordination of Zn^2+^ in the ZnF domain. Still, it was observed that several residues show chemical shift deviations or loss in intensity at lower temperature, indicating that even in the presence of Zn^2+^, temperature is influencing the local environment of the protein. These results indicate that Zn^2+^ ions protect FUS from cold denaturation and might provide a direct link between zinc dyshomeostasis and ALS pathology (see discussion).

**Figure 6.**
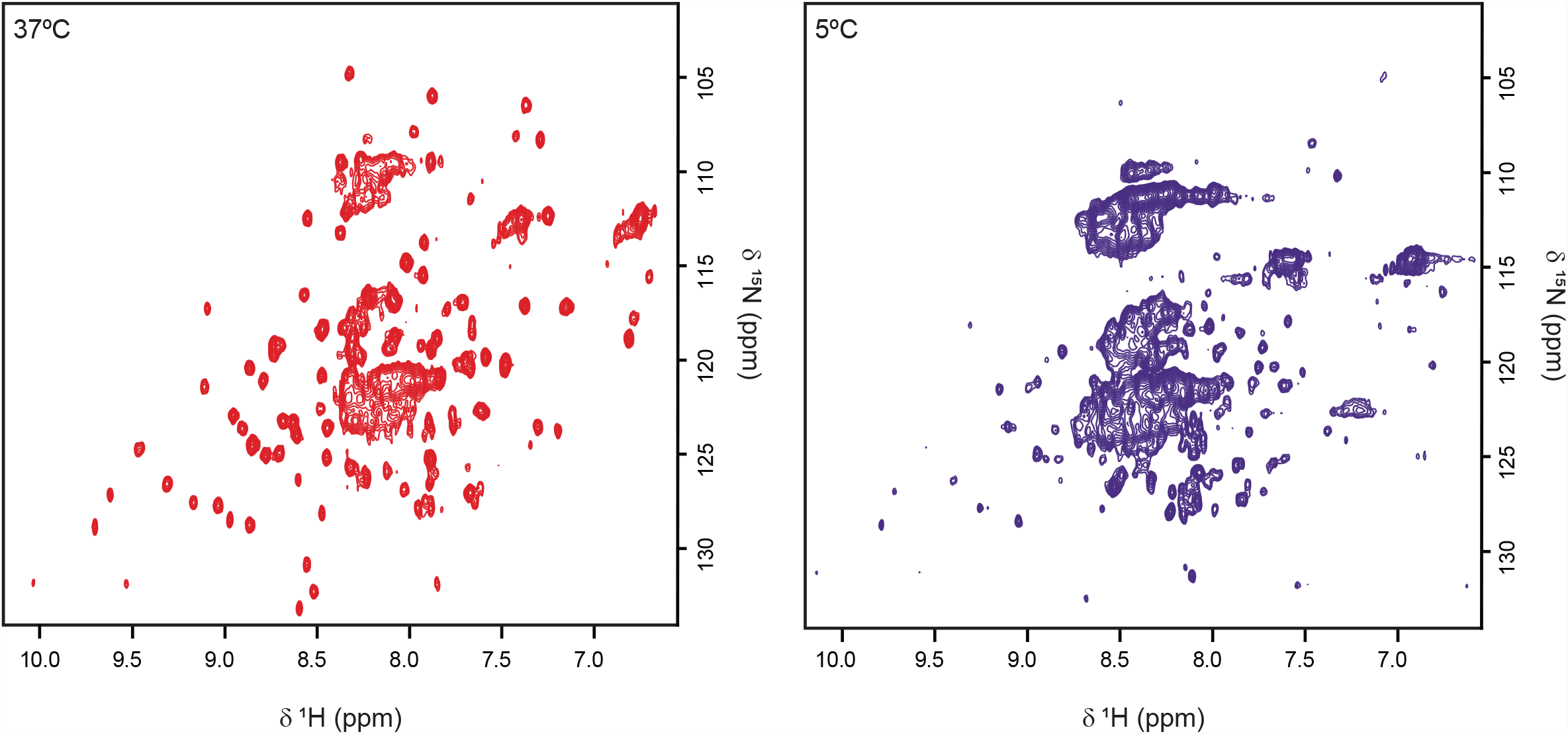
In the presence of ZnCl_2_ cold denaturation of FL FUS is not complete above 0ºC. ^1^H-^15^N TROSY-HSQC spectra of 30 μM FL FUS in the presence of 100 μM ZnCl_2_, at different temperatures, revealing that cold denaturation is not complete at 5ºC (blue) opposed to in the absence of ZnCl_2_.This suggests that proper coordination of ZnF domain is pivotal for the process to occur above 0ºC.

Taken together these results show that FUS undergoes gradual cold denaturation upon cooling towards 0ºC, at physiological pH, determined by the stability of the ZnF.

## Discussion

Studies of FL FUS are particularly challenging due to the inherent difficulty in handling FUS samples. This protein is highly prone to precipitation and non-specific binding to surfaces (Emmanouilidis *et al*, 2021). Even the simple act of pipetting can cause the protein to amorphously aggregate (Wong *et al*, 2020). Notwithstanding the difficulties, in this study we were able to establish a protocol to produce stable full-length FUS.

Our results on the influence of metabolites, pH, and metal ions, showed that the conformational stability of FL FUS LLPS is highly responsive and tightly regulated by the surrounding conditions.

Just as in the case of FUS LC, phase separation of FL FUS is greatly enhanced at 150 mM of NaCl but reduced at 300 mM (Fig. 1b), which suggests that the LLPS mechanism does not have solely an electrostatic character (Burke *et al*, 2015). We also found that FUS folded domains undergo cold denaturation, which is innately dependent on the ZnF domain stability. FUS cold denaturation might provide a pathway to increase contacts with hydrophobic residues, to allow additional intra- and inter-domain hydrophobic interactions. The hydrophobic intermolecular contacts stabilizing biomolecular condensates are reportedly the main target of 1,6-hexanediol mode of action. This aliphatic alcohol is able to disrupt phase separation by interfering with weak hydrophobic interactions (Elbaum-Garfinkle, 2019; Alberti *et al*, 2019). In the presence of 1,6-hexanediol cold denaturation of FUS was still observed (Fig. 4), hinting that 1,6-hexanediol does not interfere with the structure and folding of the protein but inhibits the phase-promoting hydrophobic interactions. Moreover, stabilizing metabolites such as glucose and glutamate hinder FUS phase separation (Fig. 1d), which is consistent with intrinsic disorder being crucial for LLPS (Murthy & Fawzi, 2020).

A recent study has demonstrated that unfolding of the FUS RRM is a key driver of condensate aging, prevented by HspB8 chaperoning (Boczek *et al*, 2021). The authors also observed that after FUS condensation multiple RRM intra- and inter-molecular contacts increased inside the condensates, suggesting RRM structural changes upon LLPS. In our study, we establish that structural changes of the RRM indeed occur via a phase-separation mediator, cold shock, which leads to the cold denaturation of the domain. This phenomenon could also help explain why FUS condensate aging occurs over time. Persistent cold shock may incite the RRM domain to remain unfolded for extended times which in turn drives condensate degeneration. In addition, we reveal that the ZnF domain also undergoes unfolding, and that the overall cold denaturation process of FUS depends on the stability of this domain. In the presence of supplemental zinc chloride, cold unfolding is significantly reduced. Zinc ions might be acting, as opposed to urea, shifting cold denaturation equilibrium towards lower temperatures by stabilizing the ZnF domain. This way, Zn^2+^ prevents complete cold denaturation of FUS at a temperature above 0ºC. The observed role of Zn^2+^ in FUS cold denaturation mechanism might be intimately linked to the zinc homeostasis dysregulation that is associated with ALS pathology (Sirabella *et al*, 2018). Other ALS-related protein, Cu/Zn-superoxide dismutase 1 (SOD1) is also regulated by Zn^2+^ ions (Rakhit & Chakrabartty, 2006). In the absence of Zn^2+^ it displays propensity to misfold and self-aggregate in toxic amyloid-like species which are found in 20% of ALS patients (Sirangelo & Iannuzzi, 2017). Without Zn^2+^, FUS might likewise display a propensity to phase-separate due to the cold denaturation mechanism which could in turn trigger aggregation. We hypothesize that FUS undergoes cold denaturation, which exposes otherwise buried hydrophobic residues, promoting LLPS via additional hydrophobic interactions, as a response to cold shock (Fig. 7). Indeed, hydrophobic interactions are known to be stabilizing in protein liquid phase states (Murthy *et al*, 2019). Nevertheless, LLPS at low temperature is not spontaneous and there is a need for certain conditions, such as salt concentration and pH. For this reason, LLPS is not observed when FUS is exposed to cold shock in non-optimal phase separating conditions or in the presence of LLPS inhibitor, 1,6-hexanediol. However, cold denaturation is still observed in these circumstances. FUS cold denaturation might promote LLPS favorable hydrophobic interactions as a mechanism preceding phase separation, but LLPS only occurs in favorable conditions. FUS structural changes with temperature may provide a fast-acting cellular response to cold stress, allowing the rapid condensate assembly and dissolution mediated by the unfolding of the globular domains.

**Figure 7.**
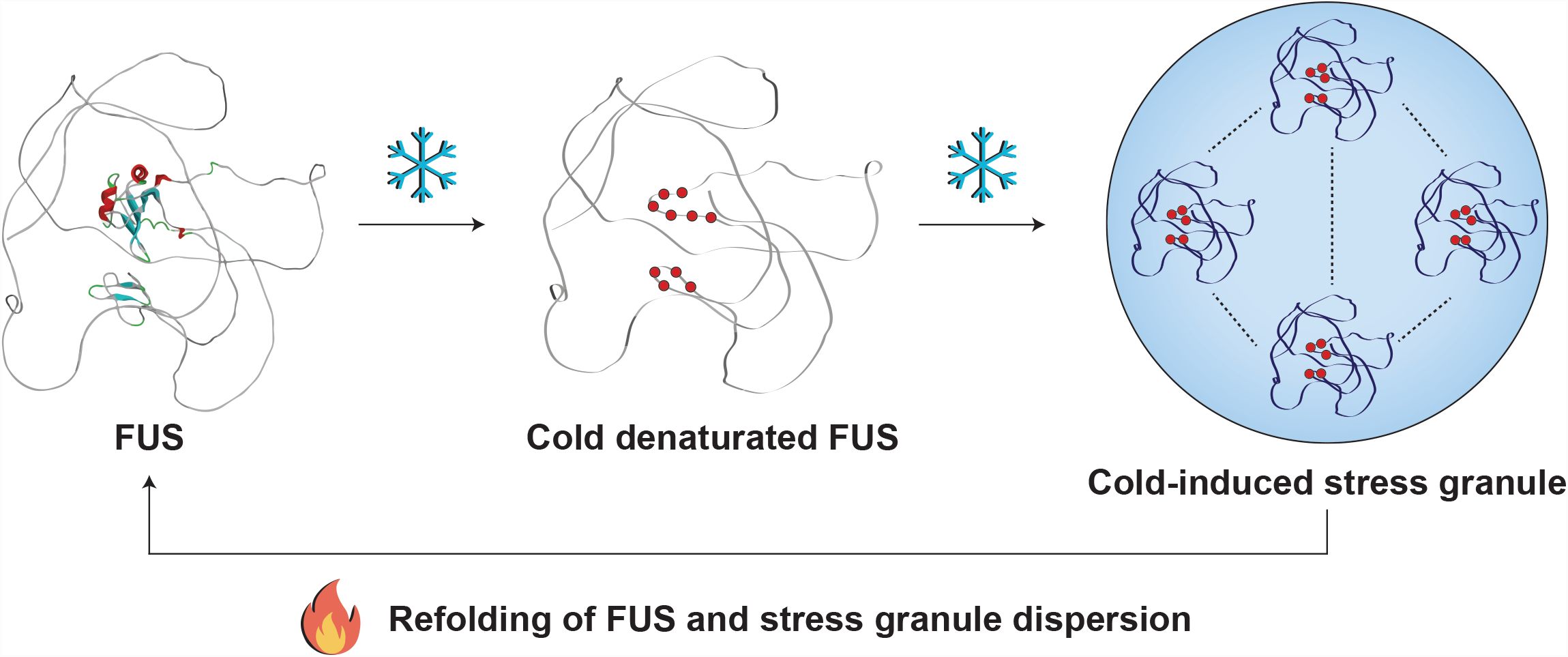
Schematic hypothesis of the implication of FUS cold denaturation on the formation of stress granules. In physiological conditions, FUS ordered domains are properly folded but upon cold stress, the domains undergo cold denaturation. This allows the exposure of buried hydrophobic residues (represented as red dots) that participate in LLPS-promoting hydrophobic interactions, leading to the assembly of stress granules. Upon temperature increase and reinstatement of physiological conditions, FUS ordered domains refold and cold-induced stress granules disassemble. FL FUS protein structure predicted by AlphaFold (Varadi *et al*, 2022; Jumper *et al*, 2021).

## Materials and Methods

### Production of FL-FUS protein

Recombinant MBP-FUS construct was kindly gifted by Nicolas Fawzi (Addgene plasmid #98651) and was expressed in BL21 (DE3) *E. coli* cells. Non-labeled FUS was expressed in LB medium supplemented with 50 μg/mL kanamycin, while uniformly ^13^C/^15^N-labeled FUS was expressed in M9 medium supplemented with 50 μg/mL kanamycin containing ^13^C-glucose and ^15^N ammonium chloride as the sole carbon and nitrogen source, respectively. Recombinant protein expression was induced at mid-exponential growth (OD_600_ 0.7-0.8) with 1 mM isopropyl-β-D-1-thiogalactoside (IPTG) for 8 hours at 30ºC. Cells pellets were resuspended in 50 mM Tris-HCl, 1 M NaCl, 10% glycerol, 1 M urea, 10 mM imidazole, 50 mM glycine, 2 mM β-mercaptoethanol, 2 mM benzamidine, 0.1 mM phenylmethylsulfonyl fluoride (PMSF) and EDTA-free cOmplete mini protease inhibitor tablet (Roche), at pH 7.40 either in the absence or presence of 100 μM ZnCl_2_. Cells were lysed through sonication and the lysate was cleared by centrifugation at 15151 *xg* for 90 minutes.

All the purification and dialysis steps were performed in temperature-controlled rooms at 25ºC, and all the tubes, buffers and columns were pre-warmed at 30ºC before use. This ensured that FUS remained stable and inhibited unspecific binding to the laboratory materials and column resins.

Affinity purifications were performed on an AKTA start prime chromatography system (Cytiva). The clarified lysate was loaded onto 2x 5 mL HisTrap FF crude columns (Cytiva) and eluted with 150 mM imidazole. MBP-FUS-containing fractions were dialyzed overnight against 20 mM Tris-HCl, 500 mM NaCl, 1 M urea, 10% glycerol, 2 mM MgSO_4_, 100 μM ZnCl_2_, 2 mM β-mercaptoethanol, 2 mM benzamidine, 0.1 mM PMSF and 0.03% NaN_3_, at pH 7.40. Benzonase nuclease (Merck) was added to the sample, in a final concentration of 15 U/mL, to remove the nucleic acids bound to FUS. To remove the Benzonase, the dialyzed sample was purified using a 5 mL HisTrap HP column (Cytiva). The MBP tag was cleaved overnight using Tobacco Etch Virus (TEV) protease (produced in house) in the aforementioned buffer lacking 2 mM MgSO_4_. The cleaved product was applied to 2x 5 mL HisTrap HP columns, where FUS eluted in the flow-through. The purified protein fractions were collected and stored at 30ºC without further concentration. Before use, freshly prepared FUS was dialyzed against the desired buffer and subsequently concentrated using Vivaspin 15 Ultrafiltration units (Sartorius). Protein purity was confirmed by SDS-PAGE and ^1^H-^15^N TROSY/HSQC spectra. The A_260nm_/A_280nm_ values obtained for pure FUS-containing samples were between 0.60-0.70, indicating a nucleic acid-free protein.

### Phase separation assays

Assessment of FUS phase separation in different conditions and in the presence of metabolites was performed by turbidity assays. Experiments were carried out by monitoring the OD at 595 nm, using a Benchmark Microplate Reader (Bio-Rad).

All assays were executed with 5 μM of pure FL FUS in 20 mM Tris-HCl pH 7.00, 20 mM CAPS pH 9.40 and 20 mM CAPS pH 11.00. Exceptions were made when the pH influenced the solubility or stability of the studied metabolites, as previously referred.

In general, metabolites or RNA solutions in variable concentrations were added to a 96-flat-bottom-well clear microplate (Corning) to a final 100 μL reaction volume. Solutions were subsequently incubated for 60 minutes at 4ºC without agitation. FUS was later added, incubated for additional 10 minutes and briefly mixed, prior to the turbidity assays. All measurements were made in triplicate.

Turbidity of FUS solutions in the presence of ZnCl_2_ was only measured at pH 7.00, since Zn^2+^ forms an insoluble hydroxide at pH 9.40 and pH 11.00.

### NMR spectroscopy

All NMR experiments were acquired in Bruker Avance III 600 MHz (^1^H) and Neo 800 MHz (^1^H) NMR spectrometers (Bruker BioSpin), equipped with cryogenic probes and Z-gradients. Chemical shifts were calibrated through indirect referencing using 50 μM sodium trimethylsilylpropanesulfonate (DSS) (Eurisotop) in all samples. All data was processed using Bruker TopSpin 4.0.6 software (Bruker BioSpin) and analyzed with CARA and PINT (Agashe & Udgaonkar, 1995; Ahlner *et al*, 2013).

For the NMR studies, ^15^N-labeled FUS samples were prepared in 90% H_2_O/10% ^2^H_2_O (Eurisotop) and transferred to 3 mm NMR tubes. Protein samples were prepared to a final concentration between 30-40 μM in 20 mM Tris-HCl, 100 mM NaCl, 5% glycerol, 2 mM β-mercaptoethanol, 0.03% NaN_3_, at pH 7.00, with or without 100 μM ZnCl_2_. To ensure that FUS was dispersed in these conditions, experiments were replicated in the presence of 4% 1,6-hexanediol (negative control). As a positive control, sample which contained 30 μM ^15^N-labeled FUS that were phase separated (as measured by turbidity) was prepared as described in the <u>Supplementary experimental procedures</u>. Samples of FUS in the presence of urea were prepared by adding urea to the protein to a final concentration of 750 mM.

Temperature influence on the structure of dispersed FUS was followed by ^1^H-^15^N TROSY-HSQC and ^1^H-^15^N HSQC spectra, recorded at 37ºC, 30ºC, 25ºC, 15ºC, 10ºC and 5ºC. The ^1^H-^15^N TROSY-HSQC spectra were acquired with 64 scans on a matrix with 2048 × 128 complex points, and a sweep width of 7812 Hz (centered at the water resonance frequency) × 2311 Hz (centered at 118 ppm), in the ^1^H and ^15^N dimensions, respectively. In turn, the ^1^H-^15^N HSQC spectra were acquired on an 800 MHz NMR spectrometer with 256 scans on a matrix with 2048 × 128 complex points, and a sweep width of 3758 Hz × 9445 Hz (centered at 116.5 ppm). 3D HNCO spectra were additionally recorded at different temperatures to confirm the proper transfer of the available assignments of the LC, RRM and ZnF domains of FUS (BMRB codes #26672, #34259 and #34258, respectively) to our spectra. On the same sample, spectra were recorded from 5ºC to 37ºC to account for the process reversibility.

### Differential interference contrast (DIC) microscopy

DIC imaging was performed using a Zeiss Axio Imager D2 microscope equipped with Zeiss 10x and 40x Plan-Neofluar objectives and Zeiss DIC EC PN 10x/40x prism sliders. Samples of 5 μM FUS in 20 mM Tris-HCl, 150 mM NaCl, 100 μM ZnCl_2_ and 2 mM β-mercaptoethanol at pH 7.40, were spotted onto glass slides, covered with coverslips and micrographed upright. Images were processed with ImageJ (Fiji) (Schindelin *et al*, 2012).

## Acknowledgements

This work was supported by Fundação para a Ciência e a Tecnologia (FCT-Portugal) for funding UCIBIO project (UIDP/04378/2020 and UIDB/04378/2020) and Associate Laboratory Institute for Health and Bioeconomy – i4HB project (LA/P/0140/2020). The authors also thank FCT-Portugal for the PhD grant attributed to SF (PD/BD/148028/2019) under the PTNMRPhD Program. JO is a recipient of a Leonardo Grant from the Spanish BBVA Foundation (BBM_TRA_0203) and a Ramón y Cajal Grant (RYC2018-026042-I funded by MCIN/AEI/10.13039/501100011033 and by “ESF Investing in your future”). JO and DVL are supported by the Spanish Grants PID-2019-109276RA-I00 and PID-2019-109306RB-I00, respectively, both funded by MCIN/AEI/10.13039/501100011033. The NMR spectrometers are part of the National NMR Facility supported by FCT-Portugal (ROTEIRO/0031/2013– PINFRA/22161/2016, co-financed by FEDER through COMPETE 2020, POCI and PORL and FCT through PIDDAC). The 800 MHz spectrometer present in the “Manuel Rico” NMR laboratory (LMR-CSIC) is a node of the Spanish Large-Scale National Facility (ICTS R-LRB-MR).

The authors would also like to acknowledge Prof. Dr. Jaime Mota and Dra. Irina Franco for the technical assistance with the microscopy experiments, Master Philip O’Toole for the aid in protein production and Dr. Aldino Viegas and Dr. David Pantoja-Uceda for the support and valuable discussions in the NMR section.

## Author contributions

JO, DVL, EC conceived, designed, and supervised all the experiments. SF developed the purification protocol. SF and JO carried out the expression and purification of FL-FUS. SF performed the phase separation turbidity assays and microscopy of FUS. SF, JO, DVL, EC performed and analyzed the NMR experiments. SF drafted the writing of the manuscript with text, figures and comments provided by all authors. All authors discussed the data analysis, critically reviewed the manuscript, and approved the final version.

## References

Agashe VR & Udgaonkar JB (1995) Thermodynamics of Denaturation of Barstar: Evidence for Cold Denaturation and Evaluation of the Interaction with Guanidine Hydrochloride. Biochemistry 34: 3286–3299

Ahlers J, Adams EM, Bader V, Pezzotti S, Winklhofer KF, Tatzelt J & Havenith M (2021) The key role of solvent in condensation: Mapping water in liquid-liquid phase-separated FUS. Biophys J 120: 1266–1275

Ahlner A, Carlsson M, Jonsson B-H & Lundström P (2013) PINT: a software for integration of peak volumes and extraction of relaxation rates. J Biomol NMR 56: 191–202

Al-Fageeh MB & Smales CM (2006) Control and regulation of the cellular responses to cold shock: the responses in yeast and mammalian systems. Biochem J 397: 247–59

Alberti S, Gladfelter A & Mittag T (2019) Considerations and Challenges in Studying Liquid-Liquid Phase Separation and Biomolecular Condensates. Cell 176: 419–434

Alberti S & Hyman AA (2021) Biomolecular condensates at the nexus of cellular stress, protein aggregation disease and ageing. Nat Rev Mol Cell Biol 22: 196–213

Arakawa T & Timasheff SN (1982) Stabilization of protein structure by sugars. Biochemistry 21: 6536–6544

Arakawa T & Timasheff SN (1984) The mechanism of action of Na glutamate, lysine HCl, and piperazine-N,N’-bis(2-ethanesulfonic acid) in the stabilization of tubulin and microtubule formation. J Biol Chem 259: 4979–4986

Banani SF, Lee HO, Hyman AA & Rosen MK (2017) Biomolecular condensates: organizers of cellular biochemistry. Nat Rev Mol Cell Biol 18: 285–298

Bentmann E, Neumann M, Tahirovic S, Rodde R, Dormann D & Haass C (2012) Requirements for stress granule recruitment of fused in sarcoma (FUS) and TAR DNA-binding protein of 43 kDa (TDP-43). J Biol Chem 287: 23079–23094

Boczek EE, Fürsch J, Niedermeier ML, Jawerth L, Jahnel M, Ruer-Gruß M, Kammer K-M, Heid P, Mediani L, Wang J, et al (2021) HspB8 prevents aberrant phase transitions of FUS by chaperoning its folded RNA-binding domain. Elife 10

Boeynaems S, Bogaert E, Kovacs D, Konijnenberg A, Timmerman E, Volkov A, Guharoy M, De Decker M, Jaspers T, Ryan VH, et al (2017) Phase Separation of C9orf72 Dipeptide Repeats Perturbs Stress Granule Dynamics. Mol Cell 65: 1044–1055

Bosco DA, Lemay N, Ko HK, Zhou H, Burke C, Kwiatkowski TJ, Sapp P, Mckenna-Yasek D, Brown RH & Hayward LJ (2010) Mutant FUS proteins that cause amyotrophic lateral sclerosis incorporate into stress granules. Hum Mol Genet 19: 4160–4175

Brangwynne CP (2013) Phase transitions and size scaling of membrane-less organelles. J Cell Biol 203: 875–881

Brangwynne CP, Eckmann CR, Courson DS, Rybarska A, Hoege C, Gharakhani J, Jülicher F & Hyman AA (2009) Germline P Granules Are Liquid Droplets That Localize by Controlled Dissolution/Condensation. Science (80-) 324: 1729–1732

Buchan JR (2014) mRNP granules. RNA Biol 11: 1019–1030

Buchan JR & Parker R (2009) Eukaryotic Stress Granules: The Ins and Outs of Translation. Mol Cell 36: 932–941

Burke KA, Janke AM, Rhine CL & Fawzi NL (2015) Residue-by-Residue View of In Vitro FUS Granules that Bind the C-Terminal Domain of RNA Polymerase II. Mol Cell 60: 231–241

Chen C, Ding X, Akram N, Xue S & Luo S-Z (2019) Fused in Sarcoma: Properties, Self-Assembly and Correlation with Neurodegenerative Diseases. Molecules 24: 1622

Courchaine EM, L. A & Neugebauer KM (2016) Droplet organelles? EMBO J 35: 1603–1612

Darling AL, Liu Y, Oldfield CJ & Uversky VN (2018) Intrinsically Disordered Proteome of Human Membrane‐Less Organelles. Proteomics 18: 1700193

Deng H, Gao K & Jankovic J (2014) The role of FUS gene variants in neurodegenerative diseases. Nat Rev Neurol 10: 337–348

Dignon GL, Zheng W, Kim YC, Best RB & Mittal J (2018) Sequence determinants of protein phase behavior from a coarse-grained model. PLOS Comput Biol 14: e1005941

Dormann D, Rodde R, Edbauer D, Bentmann E, Fischer I, Hruscha A, Than ME, Mackenzie IRA, Capell A, Schmid B, et al (2010) ALS-associated fused in sarcoma (FUS) mutations disrupt Transportin-mediated nuclear import. EMBO J 29: 2841–2857

Elbaum-Garfinkle S (2019) Matter over mind: Liquid phase separation and neurodegeneration. J Biol Chem 294: 7160–7168

Elbaum-Garfinkle S, Kim Y, Szczepaniak K, Chen CC-H, Eckmann CR, Myong S & Brangwynne CP (2015) The disordered P granule protein LAF-1 drives phase separation into droplets with tunable viscosity and dynamics. Proc Natl Acad Sci 112: 7189–7194

Emmanouilidis L, Esteban-Hofer L, Damberger FF, de Vries T, Nguyen CKX, Ibáñez LF, Mergenthal S, Klotzsch E, Yulikov M, Jeschke G, et al (2021) NMR and EPR reveal a compaction of the RNA-binding protein FUS upon droplet formation. Nat Chem Biol 17: 608–614

Gal J, Zhang J, Kwinter DM, Zhai J, Jia H, Jia J & Zhu H (2011) Nuclear localization sequence of FUS and induction of stress granules by ALS mutants. Neurobiol Aging 32: 2323.e27-2323.e40

Ganassi M, Mateju D, Bigi I, Mediani L, Poser I, Lee HO, Seguin SJ, Morelli FF, Vinet J, Leo G, et al (2016) A Surveillance Function of the HSPB8-BAG3-HSP70 Chaperone Complex Ensures Stress Granule Integrity and Dynamism. Mol Cell 63: 796–810

Hofmann S, Cherkasova V, Bankhead P, Bukau B & Stoecklin G (2012) Translation suppression promotes stress granule formation and cell survival in response to cold shock. Mol Biol Cell 23: 3786–3800

Jain S, Wheeler JR, Walters RW, Agrawal A, Barsic A & Parker R (2016) ATPase-Modulated Stress Granules Contain a Diverse Proteome and Substructure. Cell 164: 487–98

Jumper J, Evans R, Pritzel A, Green T, Figurnov M, Ronneberger O, Tunyasuvunakool K, Bates R, Žídek A, Potapenko A, et al (2021) Highly accurate protein structure prediction with AlphaFold. Nat 2021 5967873 596: 583–589

Kang J, Lim L, Lu Y & Song J (2019) A unified mechanism for LLPS of ALS/FTLD-causing FUS as well as its modulation by ATP and oligonucleic acids. PLOS Biol 17: e3000327

Kaur T, Alshareedah I, Wang W, Ngo J, Moosa M & Banerjee P (2019) Molecular Crowding Tunes Material States of Ribonucleoprotein Condensates. Biomolecules 9: 71

Kroschwald S, Maharana S & Simon A (2017) Hexanediol: a chemical probe to investigate the material properties of membrane-less compartments. Matters

Kwiatkowski TJ, Bosco DA, Leclerc AL, Tamrazian E, Vanderburg CR, Russ C, Davis A, Gilchrist J, Kasarskis EJ, Munsat T, et al (2009) Mutations in the FUS/TLS gene on chromosome 16 cause familial amyotrophic lateral sclerosis. Science 323: 1205–8

Li YR, King OD, Shorter J & Gitler AD (2013) Stress granules as crucibles of ALS pathogenesis. J Cell Biol 201: 361–372

Lin Y, Protter DSW, Rosen MK & Parker R (2015) Formation and Maturation of Phase-Separated Liquid Droplets by RNA-Binding Proteins. Mol Cell 60: 208–219

Loughlin FE, Lukavsky PJ, Kazeeva T, Reber S, Hock EM, Colombo M, Von Schroetter C, Pauli P, Cléry A, Mühlemann O, et al (2019) The Solution Structure of FUS Bound to RNA Reveals a Bipartite Mode of RNA Recognition with Both Sequence and Shape Specificity. Mol Cell 73: 490-504.e6

Mateju D, Franzmann TM, Patel A, Kopach A, Boczek EE, Maharana S, Lee HO, Carra S, Hyman AA & Alberti S (2017) An aberrant phase transition of stress granules triggered by misfolded protein and prevented by chaperone function. EMBO J 36: 1669–1687

Mitrea DM & Kriwacki RW (2016) Phase separation in biology; functional organization of a higher order. Cell Commun Signal 14: 1

Molliex A, Temirov J, Lee J, Coughlin M, Kanagaraj AP, Kim HJ, Mittag T & Taylor JP (2015) Phase Separation by Low Complexity Domains Promotes Stress Granule Assembly and Drives Pathological Fibrillization. Cell 163: 123–133

Murakami T, Qamar S, Lin JQ, Schierle GSK, Rees E, Miyashita A, Costa AR, Dodd RB, Chan FTS, Michel CH, et al (2015) ALS/FTD Mutation-Induced Phase Transition of FUS Liquid Droplets and Reversible Hydrogels into Irreversible Hydrogels Impairs RNP Granule Function. Neuron 88: 678–690

Murthy AC, Dignon GL, Kan Y, Zerze GH, Parekh SH, Mittal J & Fawzi NL (2019) Molecular interactions underlying liquid−liquid phase separation of the FUS low-complexity domain. Nat Struct Mol Biol 26

Murthy AC & Fawzi NL (2020) The (un)structural biology of biomolecular liquid-liquid phase separation using NMR spectroscopy. J Biol Chem 295: 2375–2384

Patel A, Lee HO, Jawerth L, Maharana S, Jahnel M, Hein MY, Stoynov S, Mahamid J, Saha S, Franzmann TM, et al (2015) A Liquid-to-Solid Phase Transition of the ALS Protein FUS Accelerated by Disease Mutation. Cell 162: 1066–1077

Privalov PL (1990) Cold Denaturation of Protein. Crit Rev Biochem Mol Biol 25: 281–306

Protter DSW & Parker R (2016) Principles and Properties of Stress Granules. Trends Cell Biol 26: 668–679

Psychogios N, Hau DD, Peng J, Guo AC, Mandal R, Bouatra S, Sinelnikov I, Krishnamurthy R, Eisner R, Gautam B, et al (2011) The Human Serum Metabolome. PLoS One 6: e16957

Rakhit R & Chakrabartty A (2006) Structure, folding, and misfolding of Cu,Zn superoxide dismutase in amyotrophic lateral sclerosis. Biochim Biophys Acta 1762: 1025–1037

Rogelj B, Godin K, Shaw C & Ule J (2012) The Functions of Glycine-Rich Regions in TDP-43, FUS and Related RNA-Binding Proteins. In RNA Binding Proteins pp 59–74. CRC Press

Sama RRK, Ward CL & Bosco DA (2014) Functions of FUS/TLS From DNA Repair to Stress Response: Implications for ALS. ASN Neuro 6: 1–18

Sama RRK, Ward CL, Kaushansky LJ, Lemay N, Ishigaki S, Urano F & Bosco DA (2013) FUS/TLS assembles into stress granules and is a prosurvival factor during hyperosmolar stress. J Cell Physiol 228: 2222–2231

Schindelin J, Arganda-Carreras I, Frise E, Kaynig V, Longair M, Pietzsch T, Preibisch S, Rueden C, Saalfeld S, Schmid B, et al (2012) Fiji: an open-source platform for biological-image analysis. Nat Methods 9: 676–82

Scimemi A & Beato M (2009) Determining the neurotransmitter concentration profile at active synapses. Mol Neurobiol 40: 289–306

Shin Y & Brangwynne CP (2017) Liquid phase condensation in cell physiology and disease. Science (80-) 357: 1–11

Sirabella R, Valsecchi V, Anzilotti S, Cuomo O, Vinciguerra A, Cepparulo P, Brancaccio P, Guida N, Blondeau N, Canzoniero LMT, et al (2018) Ionic homeostasis maintenance in ALS: Focus on new therapeutic targets. Front Neurosci 12: 1–14

Sirangelo I & Iannuzzi C (2017) The role of metal binding in the amyotrophic lateral sclerosis-related aggregation of copper-zinc superoxide dismutase. Molecules 22: 1429

Sun Y & Chakrabartty A (2017) Phase to Phase with TDP-43. Biochemistry 56: 809–823

Uversky VN, Kuznetsova IM, Turoverov KK & Zaslavsky B (2015) Intrinsically disordered proteins as crucial constituents of cellular aqueous two phase systems and coacervates. FEBS Lett 589: 15–22

Varadi M, Anyango S, Deshpande M, Nair S, Natassia C, Yordanova G, Yuan D, Stroe O, Wood G, Laydon A, et al (2022) AlphaFold Protein Structure Database: massively expanding the structural coverage of protein-sequence space with high-accuracy models. Nucleic Acids Res 50: D439–D444

Wang J, Choi JM, Holehouse AS, Lee HO, Zhang X, Jahnel M, Maharana S, Lemaitre R, Pozniakovsky A, Drechsel D, et al (2018) A Molecular Grammar Governing the Driving Forces for Phase Separation of Prion-like RNA Binding Proteins. Cell 174: 688-699.e16

Wong K-B, Freund SMV & Fersht AR (1996) Cold Denaturation of Barstar: 1H, 15N and 13C NMR Assignment and Characterisation of Residual Structure. J Mol Biol 259: 805–818

Wong LE, Kim TH, Muhandiram DR, Forman-Kay JD & Kay LE (2020) NMR Experiments for Studies of Dilute and Condensed Protein Phases: Application to the Phase-Separating Protein CAPRIN1. J Am Chem Soc 142: 2471–2489

Xiang S, Kato M, Wu LC, Lin Y, Ding M, Zhang Y, Yu Y & McKnight SL (2015) The LC Domain of hnRNPA2 Adopts Similar Conformations in Hydrogel Polymers, Liquid-like Droplets, and Nuclei. Cell 163: 829–839

Yang S, Warraich ST, Nicholson GA & Blair IP (2010) Fused in sarcoma/translocated in liposarcoma: A multifunctional DNA/RNA binding protein. Int J Biochem Cell Biol 42: 1408–1411

